# Stock assessment of the shorthead anchovy *Encrasicholina heteroloba* in Tanzania’s coastal waters

**DOI:** 10.64898/2026.02.18.706525

**Authors:** Changoma Marko Fransis, Takashi Fritz Matsuishi

## Abstract

The shorthead anchovy *Encrasicholina heteroloba* is a small pelagic species of significant economic and nutritional importance in Tanzania. However, reliable stock assessment remains challenging because of limited data. This study evaluates the status of *E. heteroloba* in Tanzania’s coastal waters and provides management insights for sustainable exploitation. A total of 32,324 specimens were sampled between July 2023 and December 2024. Length-frequency data were cleaned, bootstrapped, and analyzed using the ELEFAN routine to estimate von Bertalanffy growth parameters and mortality rates. Stock biomass and abundance were estimated through virtual population analysis. Estimated parameters were asymptotic length (*L*_∞_) = 8.42 cm total length, growth coefficient (*K*) = 2.08, growth performance index (∅′) = 2.19, current fishing mortality (*F*_curr_) = 0.85, exploitation rate (*E*_curr_) = 0.40, and length at first capture (*L*_curr_) = 6.10 cm total length. Natural mortality (*M*) was 1.25 and total mortality (*Z*) was 2.10. Overall biomass and stock abundance were 72,725 metric tons and 4.37 × 10^10^ individuals, respectively. Estimates of biological reference points were *F*_0.1_= 1.57, *F*_0.5_ = 1.06, and *F*_max_= 3.34. The results indicate that the stock of *E. heteroloba* in this region is currently underexploited. Fishing mortality could be moderately increased to enhance yield-per-recruit sustainably.

## Introduction

Global anchovy production has fluctuated considerably, showing an overall decline since 2010, largely driven by high fishing pressure for aquaculture feeds, fish oil, and human consumption (Barange et al. 2014; FAO 2025). Anchovies of family Engraulidae are also key forage fishes, and variability in their populations can cascade through marine ecosystems, affecting predator species and overall ecological stability (Cury et al. 2011). These pressures underscore the need for sustainable management to prevent their further depletion and to maintain ecosystem health.

In Tanzania, pelagic fisheries are predominantly multi-species, with small pelagic fish primarily in the family Engraulidae, locally known as *dagaa* (Fujimoto 2018), forming a critical component (Bodiguel and Breuil 2015). Among these, marine anchovies such as the shorthead anchovy *Encrasicholina heteroloba* are particularly abundant, as they occur extensively along the coast of Tanzania, with significant fisheries at Pangani, Mkinga, Bagamoyo, Dar es Salaam, Mafia and Kilwa (MLF 2015 Groeneveld et al. 2016; Peter et al. 2023), and are primarily harvested using ring nets (Losse 1966; Muhando and Rumisha 2008; MLF 2016).

In 2020, an estimated 8,055 metric tons (t) of marine anchovy were harvested, representing approximately 26% of the national annual fish catch; however, this figure may underestimate actual catches because informal trade in regional markets is common (MLF 2015; Ibengwe et al. 2022). Marine anchovies are a vital source of protein for Tanzanian households and contribute to regional markets as processed and sun-dried products, as well as for export (Fujimoto 2018). With over 80% of dagaa production consumed domestically (Muhando et al. 2008; Bodiguel and Breuil 2015; Fujimoto 2018), catches of marine anchovies not only support food security but also underpin local livelihoods, emphasizing the ecological and socio-economic significance of these species.

Marine anchovy fishing in the region is strongly influenced by the lunar cycle, with peak activity occurring during 15–20 moonless nights per month (Nhwani and Nhwani 1988). Fishing patterns are further shaped by seasonal monsoon winds, which move through four distinct phases: the north-east monsoon (November–February), inter-monsoon (March–April), south-east monsoon (May–August), and the second inter-monsoon (September–October). This combination of lunar and seasonal cycles enables nearly year-round anchovy harvesting (Mayala et al. 2016; Jebri et al. 2020; Sekadende et al. 2020).

Basic information on common parameters and stock status indicators such as asymptotic length (*L*_∞_), growth coefficient (*K*), stock size and biomass, biological reference points (BRPs), length at first capture, and exploitation rate (*E*) are essential for assessing a fish stock and its sustainability (von Bertalanffy 1938; Beverton and Holt 1957; Sparre and Venema 1998). The parameters *L*_∞_ and *K* help quantify whether a species is shorter-lived and faster-growing or long-lived and slower-growing, whereby a higher *K* value indicates rapid growth with a shorter lifespan, and slower growth with a longer lifespan for a lower *K* value (Pauly and Munro 1984). These parameters serve as essential inputs to yield-per-recruit (YPR) and other stock assessment models used to estimate stock biomass, stock size, current fishing mortality, *E*, and BRPs (Beverton and Holt 1957; Sparre and Venema 1998). Such information is fundamental for understanding stock status, evaluating exploitation levels, and establishing management benchmarks to ensure long-term sustainability of the stock.

However, research on stock assessment of the marine anchovy stocks in Tanzanian waters has been limited, often constrained by data scarcity. Recent research has often focused on socio-economic aspects, water quality/productivity, the biological or conservation status of the fisheries environment, cross-border trade, and value-chain enhancement (Fujimoto 2018; Kamukuru et al. 2020; Ibengwe et al. 2022, 2023). The study employed bootstrapping and analyzed the bootstrapped length-frequency data using TropFishR R package to estimate growth-curve parameters, mortality rates, exploitation rates, stock size and biomass, and BRPs from YPR analyses. Median bootstrap estimators (Fransis et al. 2026) were used to derive the final stock assessment metrics. The results will improve our understanding of marine anchovy population dynamics and establish management benchmarks for sustainable exploitation in this region.

Therefore, this study aimed to assess the stock status of *E. heteroloba*, a dominant species in the small pelagic fisheries in Tanzania’s coastal waters.

## Materials and methods

### Study sites

Three sampling sites along Tanzania’s northeastern Indian Ocean coast were selected for this study: Mkinga (4.80° S, 38.90° E), Kipumbwi (5.62° S, 38.88° E), and Bagamoyo (6.45° S, 38.90° E) (Fig. 1). The sites were chosen owing to their prominence in year-round anchovy fisheries. Additionally, the sites have stationed fisheries officers who were utilized as enumerators for on-site data collection; the enumerators were selected based on their educational background and prior experience in collecting fisheries data, particularly length and weight measurements from observed catch landings.

**Fig. 1.**
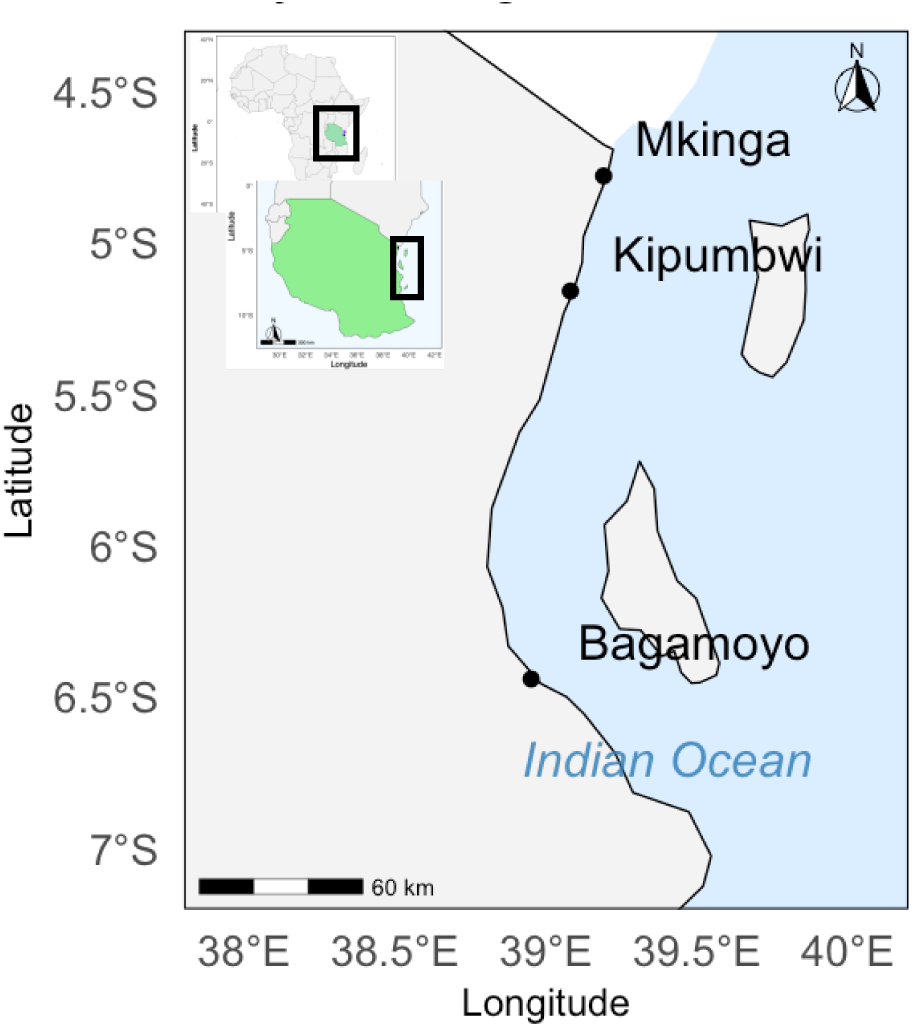
Map showing the three sites along the coast of Tanzania where *Encrasicholina heteroloba* sampling and data collection was conducted

### Sampling design and length-frequency data collection

The study focused anchovies caught using ring nets with a uniform mesh size of 8 mm. To ensure consistency, precision, and accuracy during data collection, all enumerators received on-site training on the procedures and use of provided equipment. For reliable and standardized measurements, a custom PVC measuring board was designed; PVC was selected because it is waterproof, lightweight, and durable, making it suitable for extended field use.

Fish total length (TL) was measured to the nearest 0.1 cm using a measuring board, and weight (g) to the nearest 0.1 g using a digital scale. Fish samples were randomly collected from observed landings. Sampling was conducted over seven fishing days per month at each site, though specific sampling days varied depending on the local fishery operations. Species identification followed standard identification keys. The sampling period spanned 18 months, from July 2023 to December 2024. On each sampling day, ∼100 fish were recorded per site, yielding a total of ∼2,100 samples per month.

### Structured data cleaning

To minimize the impact of outliers caused by human mistakes in parameter estimation, the dataset comprising 33,960 raw fish length measurements (cm)and individual weights (g) was first inspected for missing values, empty rows, inconsistent units, and abnormal decimal formatting, and any obvious data-entry errors were corrected. Two primary variables (length and weight) were retained in their original measurement units, and additional columns were generated to support diagnostic screening. The length–weight relationship was fitted using the non-linear model *W̃*_*i*_ = *aL*_*i*_^*b*^, and predicted weights (*W̃*) were calculated for each fish. Residuals were obtained as *d*_*i*_ = *W*_*i*_ − *W̃*_*i*_, and squared residuals were used to compute residual variance and standard deviation (SD). These values provided an objective measure of the dispersion of observations around the fitted curve.

To identify potential outliers, residuals were compared against the 95% deviation range, defined as mean ± 1.96 · SD observations, and residuals exceeding this threshold were identified as outliers and removed. Through this method, 1,636 data points (4.8% of the original dataset) were excluded, resulting in a cleaned dataset of 32,324 observations suitable for analysis. Regression analysis was then performed to determine the relationship between TL and weight of *E. heteroloba* using the length–weight relationship equation (Froese 2006; Wang et al. 2011). The quality of regression was indicated by the coefficient of determination (*R*^2^).

### Estimation of growth-curve parameters

Non-parametric bootstrapping was applied to the original *E. heteroloba* length-frequency (LFQ) data to produce pseudo-LFQ datasets (resamples), followed by ELEFAN (Pauly 1980) analysis in the TropFishR package (Mildenberger et al. 2017) to estimate growth parameters. First, a bootstrapped dataset was modified using a bin size of 0.25 cm, calculated based on the maximum length obtained in that dataset (Wang et al. 2020). The modified dataset was then restructured using a moving average (MA) of 5. Subsequently, the ELEFAN_GA function was used to estimate *L*_∞_ and *K* values for each resample. This function employs a genetic algorithm optimization technique, where parameters are treated as genes within a population (Mildenberger et al. 2017).

The search space for initial values was defined as ±20% of the nominal values, resulting in upper and lower limits for *L*_∞_ of 10.0 cm and for *K* of 2.0 year^−1^. The von Bertalanffy growth function (von Bertalanffy 1938) was used to estimate von Ber talanffy growth curve (VBGC) parameters by fitting the model to the LFQ data. The estimated VBGC parameters were used to calculate the growth performance index (∅′) as expressed by Pauly and Munro (1984). Finally, median bootstrap resampling estimators were used to derive assessment metrics for each stock parameter.

### Estimation of exploitation status

The linearized length-converted catch curve (LCCC) method, as defined below (Pauly 1990; Pauly et al. 1995), was used to estimate the instantaneous total mortality rate (*Z*). Natural mortality (*M*) was estimated by using the empirical formula of Tanaka (1960). Fishing mortality (*F*) was estimated based on the relationship *F* = *Z* − *M*. The exploitation rate (*E*) was calculated as *E* = *F*/*Z* (Gulland 1971).

### Stock-size estimation and YPR analysis

The virtual population analysis (VPA) method of Jones (1984) was used to reconstruct the standing stock biomass and stock size, as well as to estimate the fishing mortality of each length class using the VBGC parameters estimated by ELEFAN.

The fishing mortality derived from the LCCC was used to estimate the fishing mortality of the last length class, called terminal *F*. Last length classes with low catch numbers were grouped into plus-groups.

The Jones’ length-based calculations used two key equations:

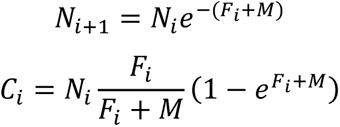

Where *i* denotes the length-class index, *N*_*i*_ is the stock size in numbers, *C*_*i*_ is the catch representing stock biomass, *F*_*i*_ is the fishing mortality, and *M* is the natural mortality.

The fishing mortality-based reference points (*F*_0.1_, *F*_0.5_, and *F*_max_) for each bootstrapped LFQ dataset were predicted using the Thompson and Bell (1934) model implemented within the ELEFAN framework. The YPR model incorporates the Thompson and Bell model to estimate yield and biomass of the stock, and uses *F*, length–weight relationship parameters (*a* and *b*), and gear selectivity parameters to predict *F*_0.1_, *F*_0.5_, and *F*_max_. YPR builds on the output of the length-based VPA, *M*, and gear selectivity (Thompson and Bell 1934; Sparre and Venema 1998).

## Results

### LFQ analysis

A total of 32,324 LFQ samples of *E. heteroloba* were analyzed, with the individual lengths in the range of 3.9–10.5 cm TL. Fish weight was strongly predicted by length. The correlation between weight and length suggests that the *E. heteroloba* stock exhibited positive allometric growth (Fig. 2).

**Fig. 2.**
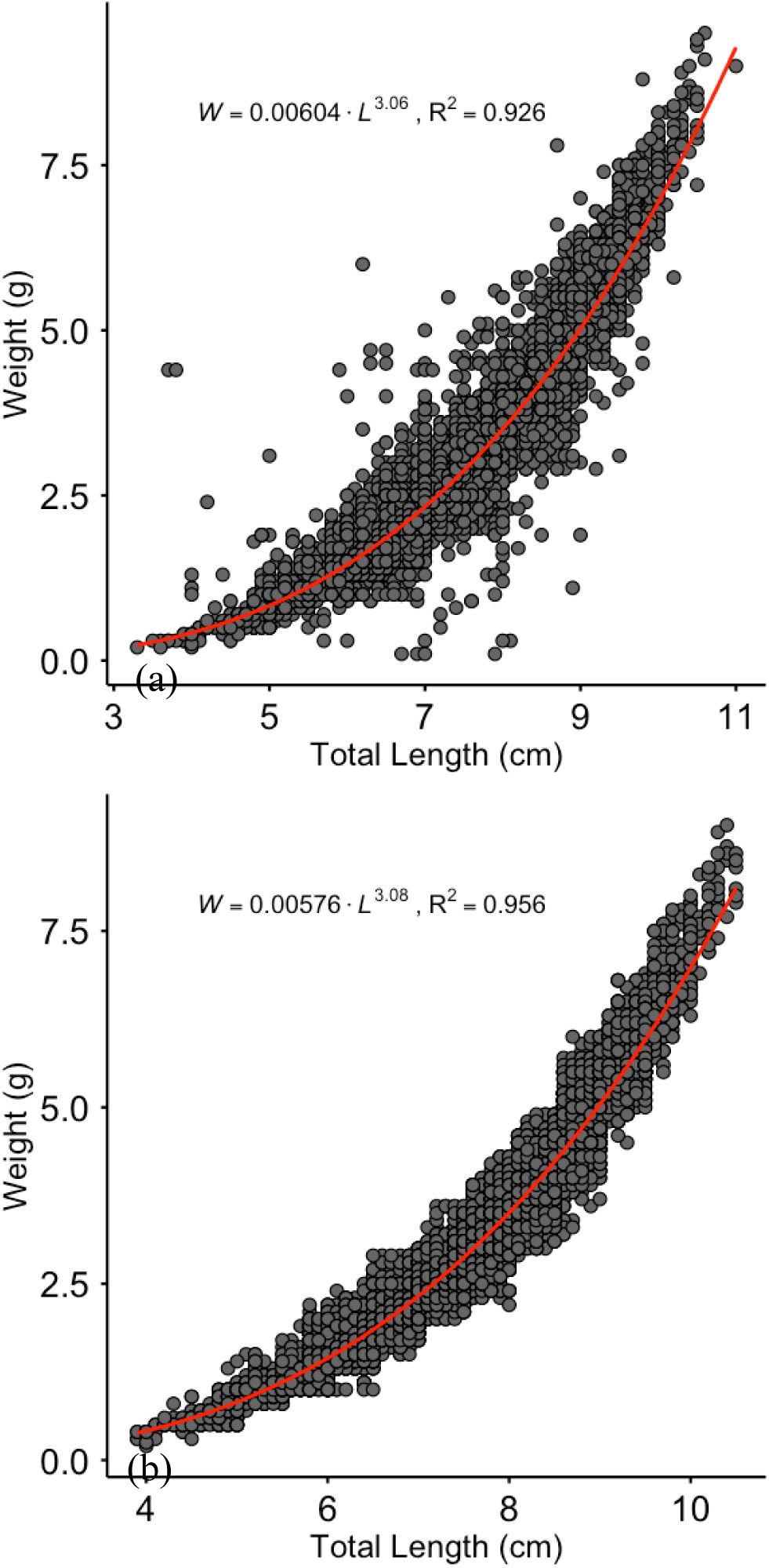
The length–weight relationship of *Encrasicholina heteroloba* (a) before (*n* = 33,960 raw length observations) and (b) after (*n* = 32,324 observations) structured data cleaning

Furthermore, the 25th–75th percentile range of the length-frequency distribution indicated a relatively wide size range within the catch, reflecting the presence of both juveniles and larger individuals. Although fish of various size classes were recorded, fish sized 6–9-cm TL were dominant, representing the modal group in the sample (Table 1, Figs. 3 and 4).

**Fig. 3.**
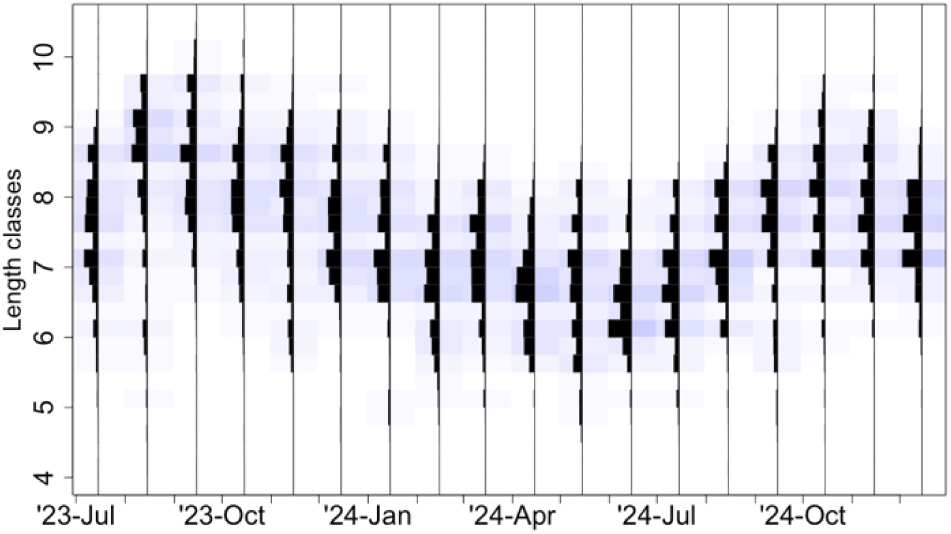
Monthly distribution and fluctuation in length classes of *Encrasicholina heteroloba* in the catches sampled at three sites on the coast of Tanzania

**Fig. 4.**
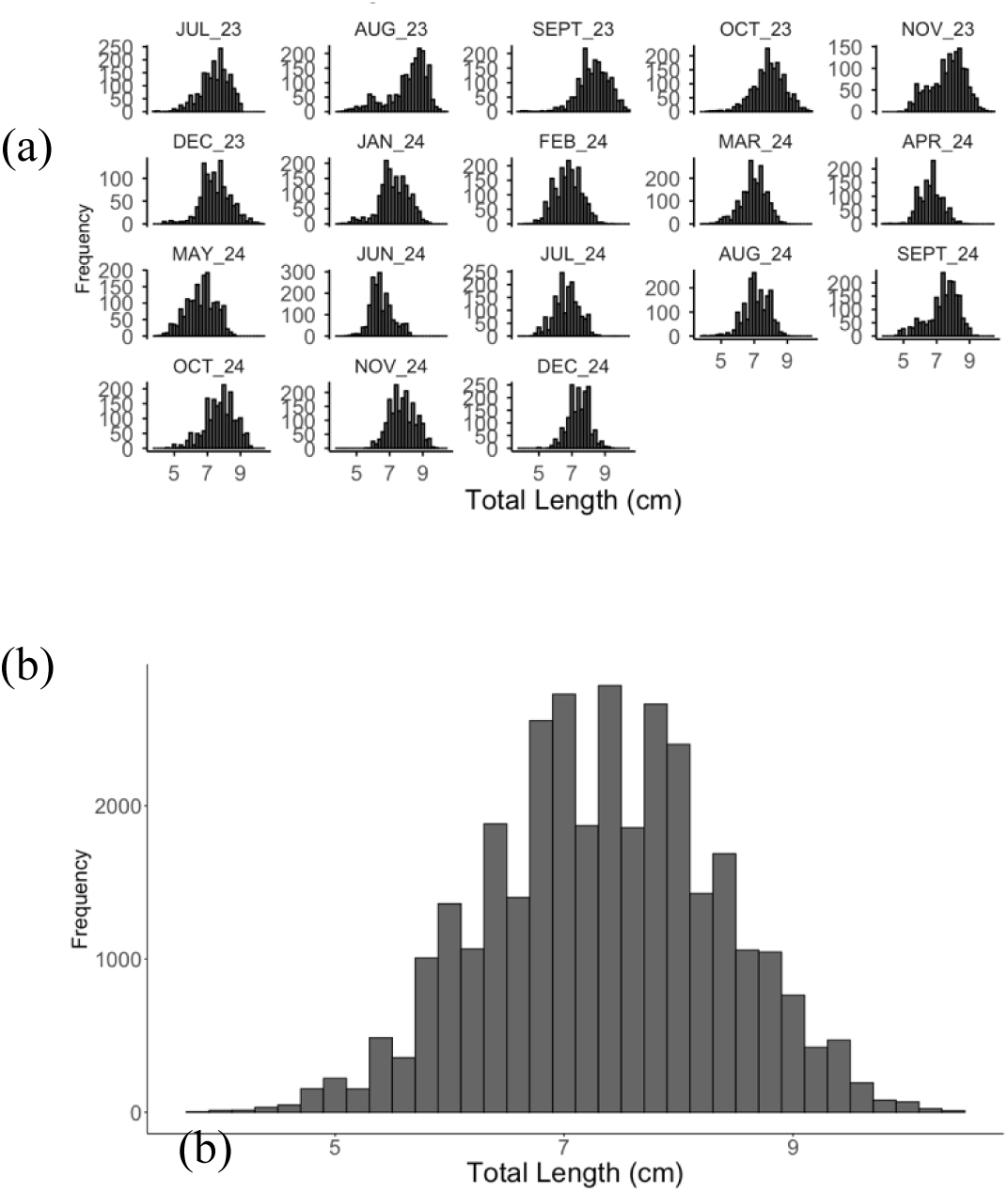
Length-frequency distributions of *Encrasicholina heteroloba* in (a) the monthly data andn(b) pooled data

**Table 1.** Results of the length-frequency analysis of *Encrasicholina heteroloba* individuals collected at three key landing sites on the coast of Tanzania.

| Sample size | Range (cm) | $R^2$ | 25th percentile | 75th percentile |
| --- | --- | --- | --- | --- |
| 32,324 | 3.9–10.5 | 0.955 | 6.7 | 8.0 |

### VBGC parameters

The estimated VBGC parameters and their 95% confidence intervals (CIs), obtained using median bootstrap estimators based on 100 bootstrap resamples, are presented in Table 2. The estimated CIs indicate overall uncertainty in the parameter estimates, with the upper CI bounds showing relatively smaller deviations from the estimated values.

**Table 2.**
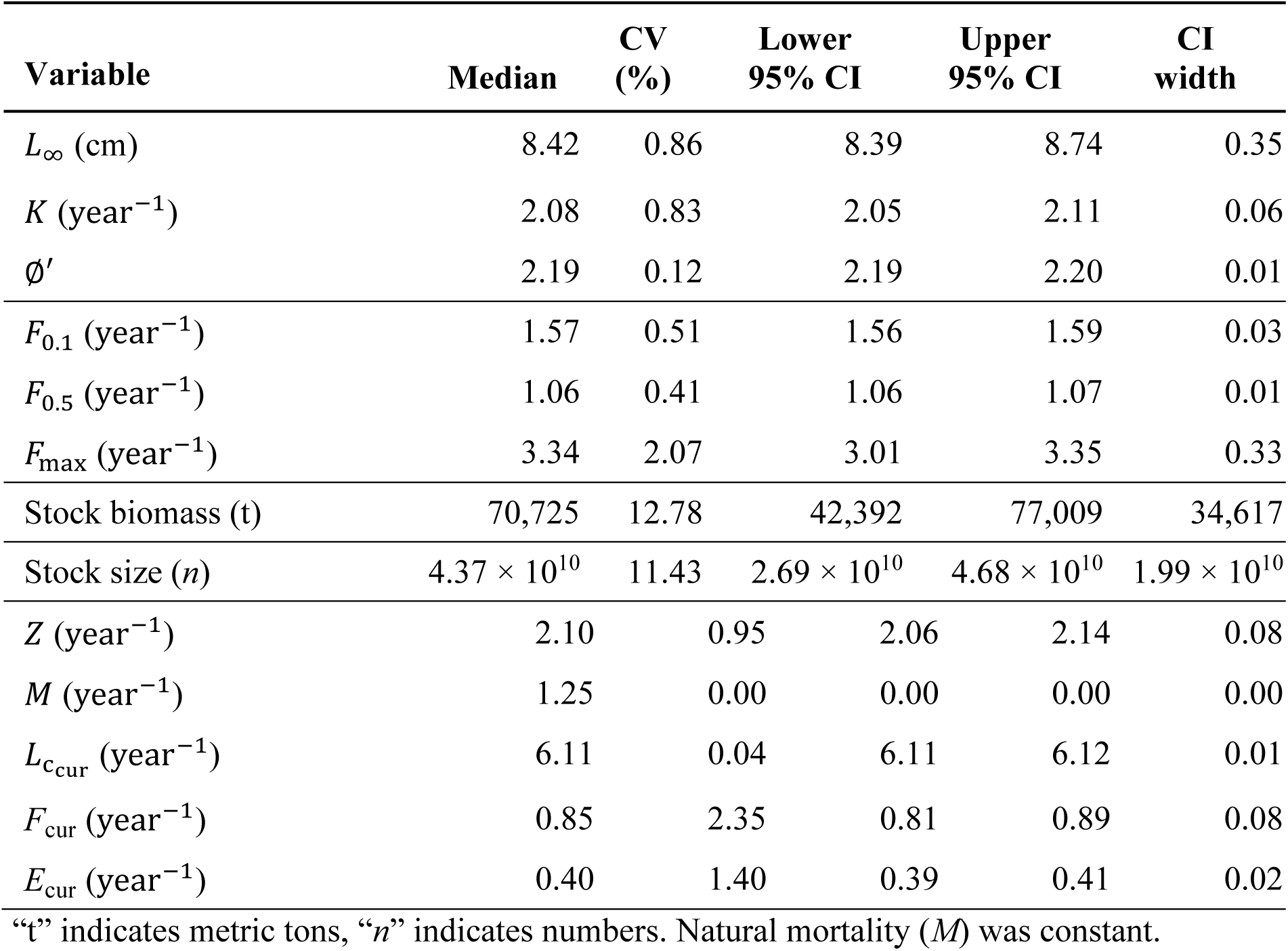
Estimated von Bertalanffy growth parameters, biological reference points, stock biomass and population abundance, current fishing mortality, and current exploitation rate of *Encrasicholina heteroloba* in the coastal waters of Tanzania

### Fishing mortality and exploitation rates

The fishing mortality and exploitation rates of the *E. heteroloba* stock are presented in Table 2. The estimated length at first capture (*L*_curr_) falls within the mid-range of the observed size distributions. Similarly, the current fishing mortality (*F*_curr_) was lower than the natural mortality (*M* = 1.25). The exploitation rate (*E*_curr_) was below the conventional optimum level of *E* = 0.5 (Gulland 1971). All parameter estimates had narrow CIs, indicating narrow uncertainty.

### Stock size and YPR output

The estimated stock biomass, derived from cohort analysis using VPA, is presented in Table 2. The VPA output indicated a gradual decline in stock numbers with increasing length, consistent with the expected mortality pattern of a short-lived species and the observed gear selectivity (Fig. 4). Fish in the size classes 5–7 cm TL were most vulnerable to fishing mortality, whereas individuals sized <5 cm TL were less affected.

The BRPs of fishing mortality estimated from the model are presented in Table 2. The graphical outputs of the YPR and biomass-per-recruit models, illustrating the current stock status (*F*_curr_ and *L*_curr_), are illustrated in Fig. 5. The estimate of *F*_curr_ was lower than all three BRPs (*F*_0.1_, *F*_0.5_, and *F*_max_). These results show that the fishing pressure exerted on the *E. heteroloba* population was relatively low to account for optimal yield.

**Fig. 5.**
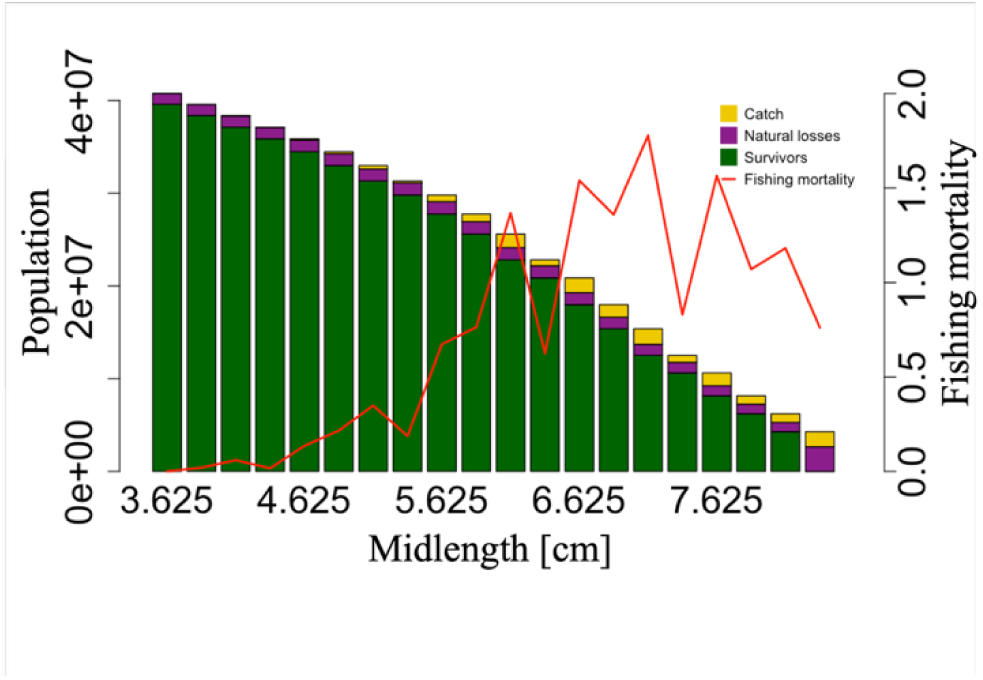
Virtual population analysis of the stock of *Encrasicholina heteroloba* in the coastal waters of Tanzania

The estimated YPR at the current levels of *F* was relatively low. The model predicted that the maximum potential yield could be obtained at *F*_max_, whereas 50% of the stock biomass could be obtained at *F*_0.5_. The model further indicated that the biologically optimal yield could be achieved when fishing mortality corresponds to *F*_0.1_. Gear-related analysis suggested that, under the present gear mesh size, the exploitation rate could be increased to produce >0.6 g per recruit without adversely affecting the spawning stock biomass (Fig. 6&7).

**Fig. 6.**
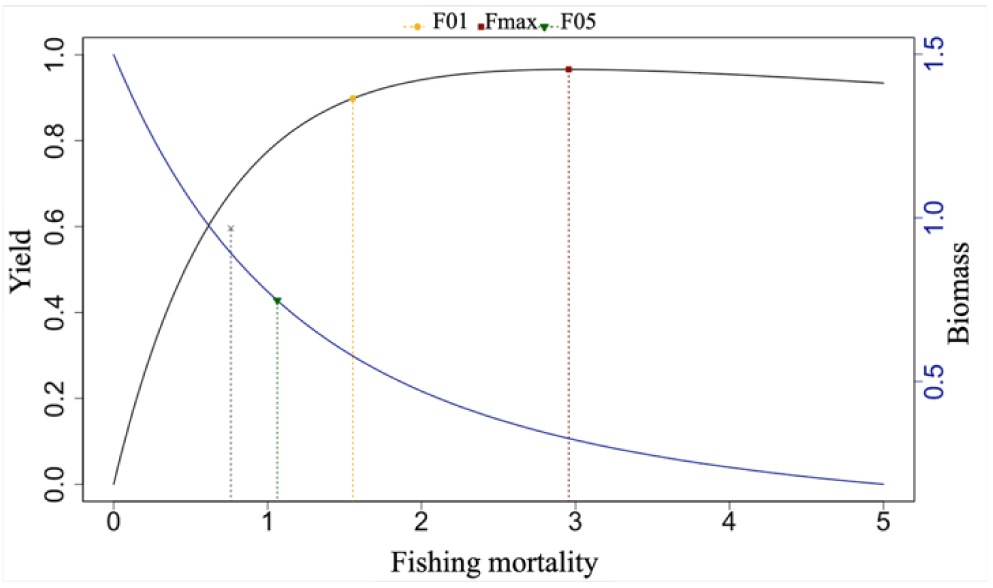
Yield-per-recruit plot showing yield, current fishing mortality (denoted by ‘x’), biological reference points (BRPs) (i. e. *F*_0.1_, *F*_0.5_, *F*_max_), and biomass per recruit for the stock of *Encrasicholina heteroloba* in Tanzania’s coastal waters. Dotted vertical lines represent current fishing mortality and the respective BRPs

**Fig. 7.**
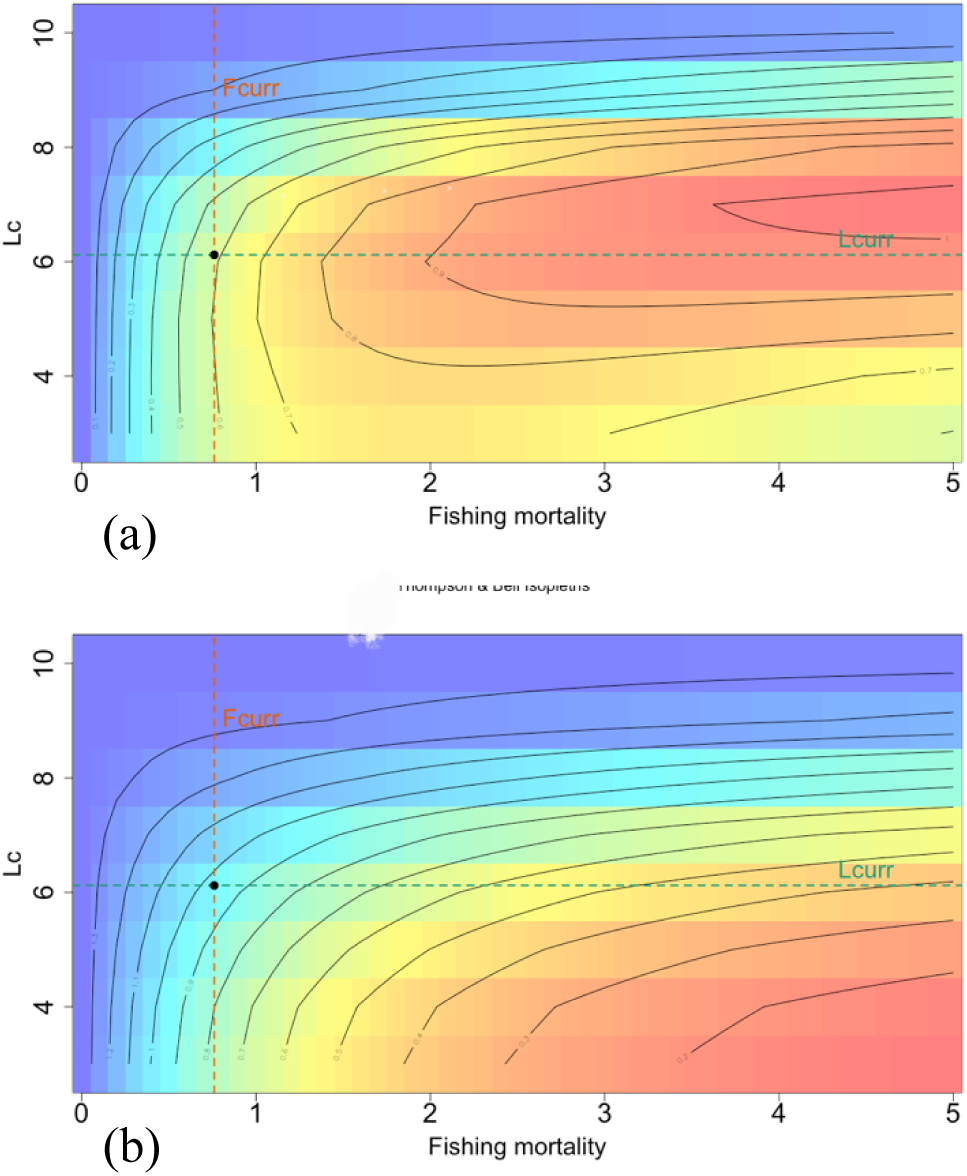
Results of the yield-per-recruit (YPR) model showing (a) the current fishing mortality (*F*_curr_) and current length at first capture (*L*_*c*urr_), alongside with the corresponding YPR and (b) biomass per recruit, for the stock of *Encrasicholina heteroloba* in Tanzania’s coastal waters. The *x*-axis represents the fishing mortality rate applied to fully exploited length classes, while the *y*-axis shows the currently estimated total length at first capture

## Discussion

In developing countries such as Tanzania, small-scale fisheries do not play a significant role in the national economic performance, despite their substantial contribution to income generation, employment, and food security among both coastal and inland freshwater fishing communities. Consequently, the sector is characterized by a limited quantity and quality of available data for effective assessment of the fishery resources, posing a major challenge for managers who must make decisions under considerable uncertainty.

In response to these data limitations, this study sought to provide the best possible assessment of the target species exploited by small scale fishers in Tanzania, thereby establishing a basis for informed management decisions. In the following paragraphs, the outcomes of the assessment routines and their implications for fisheries management are discussed.

### Length-frequency distribution

The length-frequency analysis revealed that the *E. heteroloba* catch comprised individuals of mixed length classes, with fish measuring 6–9 cm TL dominating. This suggests that the majority of analyzed *E. heteroloba* individuals belonged to mature and larger size classes. The observed mixture of length size distributions could be attributed to fishing gear selectivity, as well as to temporal and spatial variations in the sampling locations (Herrón et al. 2018).

This is supported by the 25th percentile length-frequency distributions that is within the length at first maturity for this species. This indicates that a substantial portion of individuals have reached reproductive maturity before capture (Table 1). Most mature individuals likely contribute to spawning before being removed from the population, suggesting a lower risk of recruitment overfishing (Froese 2004; Cope and Punt 2009).

Additionally, the 75th percentile length-frequency distributions surpassed the optimal length at capture, implying that a significant fraction of individuals are harvested beyond the size that maximizes their biomass contribution to the fishery (Froese et al. 2016). While this can be beneficial for maximizing yield, excessive capture of fish beyond the optimal size may indicate underexploitation or suboptimal utilization of the stock. Hence, these percentile length-frequency distributions provide insight into fishing pressure and population status of Tanzania’s exploited stock of *E. heteroloba* (Hordyk et al. 2015; Prince et al. 2015).

Monitoring shifts in length structure is crucial, as declining median lengths or shifts in percentile values toward smaller sizes often signal increased fishing pressure and the potential risk of overfishing (Beverton and Holt 1957). Thus, the observed length distribution suggests that the current exploitation pattern allows for reproductive success while capturing individuals at an appropriate size, suggesting a balance fundamental to a sustainable harvest. This pattern may also indicate continuous recruitment and overlapping generations within the *E. heteroloba* population, reflecting a stable population structure under the prevailing fishing conditions.

### Growth parameters of *E. heteroloba*

The growth of *E. heteroloba* in this study was described by von Bertalanffy growth model yielding VBGC parameters. The median *L*_∞_ (8.42 cm TL) estimated for *E. heteroloba* in this study falls with the range of previously reported values at 7.4–13.0 cm TL (Table 3). Despite this, VBGC parameters for this species remain poorly studied and relevant literature is scarce. Most available studies originate from Indonesia and the Philippines (Pauly 1978; Supeni and Dobo 2017; Juliani et al. 2019; Musel et al. 2022).

**Table 3.**
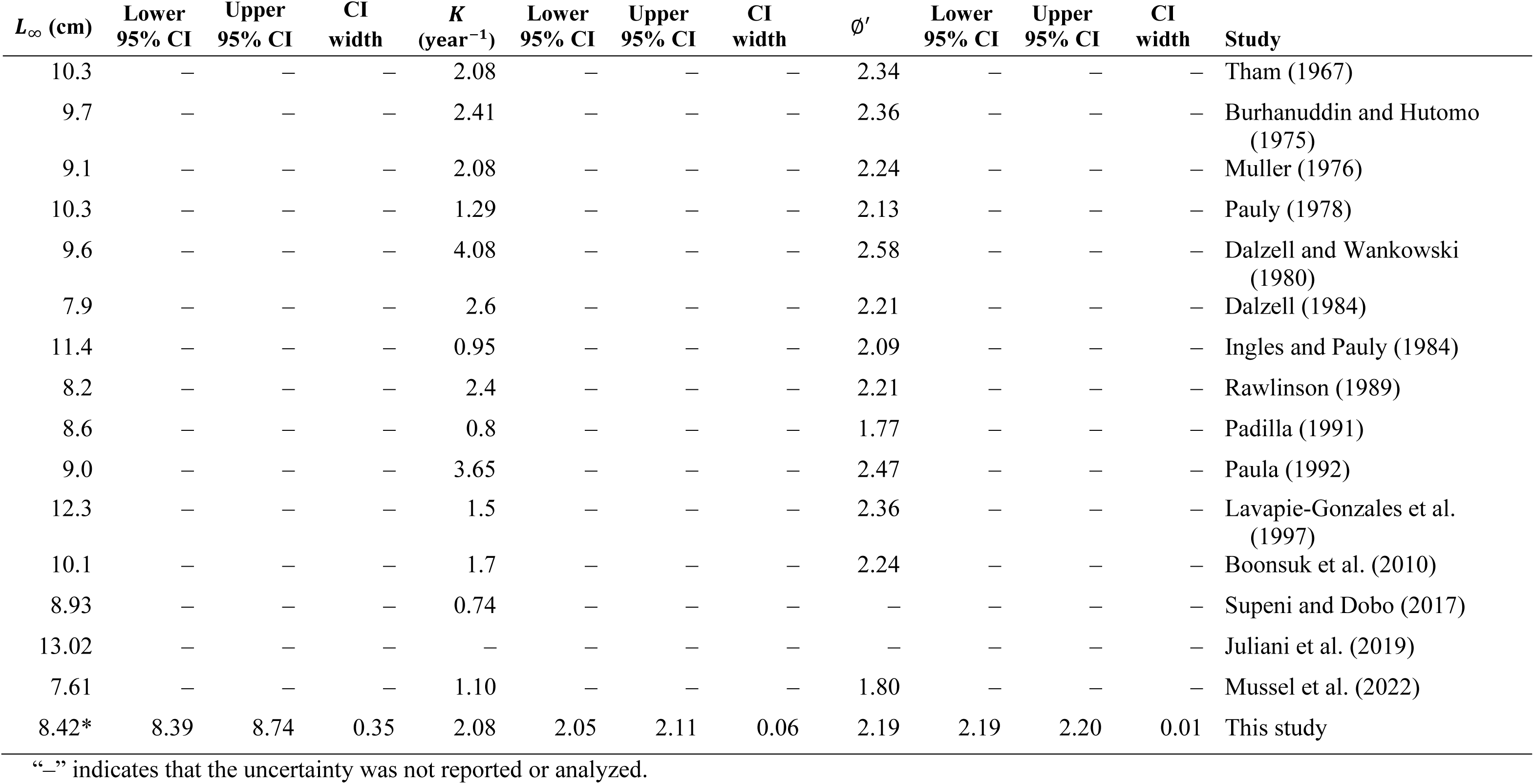
Estimated von Bertalanffy growth parameters (asymptotic length *L*_∞_, growth coefficient *K*, and growth performance index ∅′, with confidence intervals [CIs]) for *Encrasicholina heteroloba*, based on length-frequency catch data from different studies

The wide variation in reported *L*_∞_ values highlights substantial variability in growth estimates, suggesting the need for more-robust and standardized estimation approaches. This variability may, in part, be attributed to differences in sample sizes, the size distributions of sampled individuals, and length measurement methods, since TL, standard length, and fork length have all been used across studies (Dalzell 1984; Boonsuk et al. 2010; Taufani and Matsuishi 2024). The estimated median *K* (2.08) in this study is broadly consistent with previously reported values in the range of 0.48–4.45 year^−1^for similar species.

Of the two parameters, *K* is generally more variable than *L*_∞_ across studies, especially for data-limited fisheries. This is likely owing to its higher sensitivity to sampling structure, model assumptions, and estimation methods, especially when larger individuals are underrepresented in the data. Moreover, the variability in the VBGC parameter estimates might be variously attributed to the influence of climate change or differences in geographical location, diet and exploitation rate (He et al. 2016). Additionally, the relatively high *K* value and corresponding *L*_∞_ indicate that *E. heteroloba* in Tanzania’s coastal waters exhibits rapid growth and a shorter lifespan, consistent with the characteristics of a fast-growing species (Raymond et al. 2004).

The median ∅′(2.19) estimated in this study is within the range of previously reported values at 1.79–2.6 (Dalzell 1984; Boonsuk et al. 2010; Musel et al. 2022). Since ∅′ is a composite index reflecting both *K* and *L*_∞_, discrepancies among populations may suggest considerable differences in growth performance, possibly driven by regional environmental factors, ecological conditions, or inconsistency in the accuracy of the estimated VBGC parameters (Pauly and Munro 1984).

### Current fishing mortality and exploitation rate

Comparative assessment of current fishing mortality (*F*_curr_), exploitation rate (*E*_curr_) and length at first capture (*L*_curr_) is indispensable for evaluating the status of fish stocks and developing effective fisheries management strategies to ensure sustainable fishing practices. In this study, *F*_curr_ was found to be lower than the predicted BRPs of *F*_0.1_, *F*_0.5_, and *F*_max_ (Table 2), suggesting that the *E. heteroloba* stock is currently underfished.

Similarly, *E*_curr_ was below the conventional optimum harvest rate (*E* = 0.5) proposed by Gulland (1971), indicating low fishing pressure relative to stock productivity. The value of *E*_curr_ here also implies that a relatively small portion of the available *E. heteroloba* biomass is harvested annually, which is consistent with the observed low YPR and the high available biomass per recruit.

The value of *L*_curr_ further indicates that *F*_curr_ predominantly targets mature individuals, suggesting minimal risk of recruitment or growth overfishing of the *E. heteroloba* stock. The *L*_curr_ is approximately equal to the reported mean length at first maturity (*L*_50_), implying that individuals in the catch are harvested at or just after reaching sexual maturity (Fabiani et al. 2022; Froese and Pauly 2025). This observation is supported by the catch composition, which consists of a mixture of individuals from the *L*_curr_ and larger size classes, suggesting that the current exploitation pattern is likely sustainable under the existing level of fishing pressure.

Furthermore, the current gear selectivity suggests that the applied mesh size of 8 cm has no impact on immature individuals of *E. heteroloba*. However, any increase in *F*_curr_ and a shift in selectivity toward smaller sizes could elevate the risk of growth or recruitment overfishing. Therefore, maintaining or slightly increasing *L*_curr_ is recommended as a precautionary measure.

### Stock status

The VPA estimates for *E. heteroloba* indicate that losses in the stock are mainly attributable to natural mortality for fish up to 5 cm TL, but that individuals become increasingly vulnerable to the fishing gear at sizes greater than 5 cm. However, the vulnerability of the fully exploited size classes to the fishing gear remains subordinate to natural causes, indicating that natural causes contribute slightly more than fishing to total mortality (Fig. 5). This is supported by the current estimates of biomass (70,725 t) and population abundance (4.37 × 10¹⁰ individuals), which together suggest that the *E. heteroloba* stock is stable. Furthermore, the observed stock stability can be attributed to the characteristic productivity of tropical fisheries, which is supported by continuous reproduction and recruitment throughout the year (Matsuishi 2025).

The *F*_curr_ of *E. heteroloba* was lower than the BRPs, which aligns with relatively low exploitation rate reported here, suggesting an underexploited stock status. The prediction plots indicate that yield could be increased by raising fishing mortality, without the need to adjust *L*_curr_ or modify the current fishing gear mesh size (Fig. 6). Specifically, adjustments toward the BRPs benchmarks of *F*_0.1_ and *F*_0.5_(excluding *F*_max_) could enhance yield without compromising sustainability of the fishery (Herrón et al. 2018).

Moreover, applying *F*_max_ as the maximum fishing mortality that produces maximum sustainable yield (MSY) should be interpreted with caution. This is because *F*_max_ can be overestimated, potentially leading to fishery collapse if used as a benchmark for achieving MSY. Relying solely on *F*_max_ as a reference point for fisheries management has been criticized in several studies, as it often exceeds the fishing mortality at MSY (Mace 2001).

Similarly, Thordarson et al. (2006) pointed out that many stocks exhibit YPR curves that are asymptotic or flat-topped, making it easy to exceed *F*_max_ and thus increase the risk of overfishing. Furthermore, estimation of this BRP does not account for other factors, such as spawning-stock biomass levels, population structure, and density-dependent growth effects, all of which can lead to overestimation of MSY and potentially result in overfishing or stock collapse (Horbowy and Hommik 2022).

The yield isopleth diagrams indicate that increasing fishing mortality at the same *L*_curr_ will increase yield (Fig. 7). Similarly, this increase in fishing mortality could reduce the biomass per recruit to within sustainable limits. These results suggest that a moderate increase in fishing effort could optimize harvest from the *E. heteroloba* fishery in this region. Such adjustments could achieve both sustainable exploitation and improved socioeconomic returns (Harlyan et al. 2019).

## Conclusions

The findings of this study indicate that the stock of *E. heteroloba* in Tanzania’s coastal waters is currently underexploited, as evidenced by our estimates of stock biomass, population abundance, a low exploitation rate, and relatively low fishing mortality. Furthermore, the estimated biological reference points, growth parameters, and stock biomass information provide insights that can support evidence-based fisheries management and promote the sustainable utilization of *E. heteroloba* in this region. The model results also suggest that the current fishing mortality could be increased moderately (toward the estimated BRPs) to enhance yield-per-recruit without compromising sustainability.

As a tropical small pelagic species, *E. heteroloba* presents unique challenges for traditional stock assessment methods because of limited fishery data and based on its continuous spawning behavior and an absence of distinct seasonal cohorts, unlike temperate species. These biological characteristics complicate cohort tracking and reduce the effectiveness of conventional length-based stock assessment approaches. In this context, the application of the median bootstrap estimator approach using a length-frequency dataset proved particularly advantageous, offering a robust framework to estimate population parameters under data-poor conditions.

## Acknowledgements

The authors acknowledge financial support from the Government of the United Republic of Tanzania, through the Ministry of Education, Science and Technology, under the Higher Education for Economic Transformation (HEET) Project, which funded the PhD scholarship of Changoma Marko Fransis and fully supported this study. Cynthia Kulongowski from Edanz (https://jp.edanz.com/ac) edited the language of a draft of this manuscript.

## Author contributions

**Changoma Marko Fransis:** Conceptualization, Methodology, Data Curation, Formal Analysis, and Investigation, Writing – original draft preparation, review and editing, Resources; **Takashi Fritz Matsuishi:** Conceptualization, Supervision and Project Administration, Methodology, Writing – review and editing.

## Data availability

The length-frequency data supporting the findings of this study were collected in the field for research purposes and are available from the corresponding author upon reasonable request.

## Declarations

## References

Barange M, Merino G, Blanchard JL, Scholtens J, Harle J, Allison EH, Allen JI, Holt J, Jennings S (2014) Impacts of climate change on marine ecosystem production in societies dependent on fisheries. Nat. Clim. Change 4:211–16. 10.1038/nclimate2119

Beverton RJH, Holt SJ (1957) On the dynamics of exploited fish populations. Fish. Invest. Ser. 2 Mar. Fish. G.B. Minist. Agric. Fish. Food 19: 533.

Bodiguel C, Breuil C (2015) Report of the meeting on marine small pelagic fishery in the United Republic of Tanzania. SFFAO/2015/34, IOC-SmartFish Programme of the Indian Ocean Commission. FAO. Ebene, Mauritius. 90.

Boonsuk S, Sumontha M, Sa-nga-ngam C, Tes-a-sen K (2010) Stock assessment of anchovies (Encrasicholina devisi (Whitley, 1940), E. punctifer Fowler, 1938 and E. heteroloba (Ruppell, 1837)) along the Andaman Sea Coast of Thailand. Tech. Pap. No. 17/2010. Marine Fisheries Research and Development Bureau, Department of Fisheries, Ministry of Agriculture and Cooperatives. 40.

Burhanuddin SM, Hutomo M (1975) A preliminary study on the growth and food of Stolephorus spp. From Jakarta Bay. Mar. Res. Indones. 14: 1 – 29.

Cope JM, Punt AE (2009) Length-based reference points for data-limited situations: applications and restrictions. Mar. Coast. Fish. 1(1): 169–186. 10.1577/C08-025.1

Cury PM, Boyd IL, Bonhommeau S, Anker-Nilssen T, Crawford RJM, Furness RW, Mills JA, Murphy EJ, Österblom H, Paleczny M, Piatt JF, Roux JP, Shannon L, Sydeman WJ (2011) Global seabird response to forage fish depletion: one-third for the birds. Science 334:1703–6. 10.1126/science.1212928

Dalzell P (1984) The population biology and management of baitfish in Papua New Guinea waters. Papua New Guinea Dept. Primary Industries Res. Rep. 84-05. 59.

Dalzell PJ, Wankowski JW (1980) The biology, population dynamics and fisheries dynamics of exploited stocks of three baitfish species: *Stolephorus heterolobus*, *S. devisi* and *Spratelloides gracilis* in Ysabel Passage, New Ireland Province. Res. Bull. Dept. Primary Ind. (Papua New Guinea) 22: 124.

Fabiani G, Chauka J, Muhando CA (2022) Population Characteristics of selected small pelagic fish species along the Tanzanian coast. Tanz. J. Sci. 48(3), 585–595. 10.4314/tjs.v48i3.6

FAO (2025) *Engraulis ringens* Jenyns,1842. In: Fisheries and Aquaculture [accessed October 21, 2025]. www.fao.org/fishery/en/aqspecies/2917/en

Froese R (2004) Keep it simple: three indicators to deal with overfishing. Fish Fish. 5: 86–91. 10.1111/j.1467-2979.2004.00144.x

Fransis CM, Kanno H, Matsuishi TF (2026) Consistent ELEFAN estimation of growth curve parameters and biological reference points using medians. AACL Bioflux 19(1): 1090–1105 (in press)

Froese R (2006) Cube law, condition factor and weight–length relationships: history, meta-analysis and recommendations. J. Appl. Ichthyol. 22: 241–253. 10.1111/j.1439-0426.2006.00805.x

Froese R, Winker H, Gascuel D, Sumaila UR, Pauly D (2016) Minimizing the impact of fishing. Fish. 17: 785–802. 10.1111/faf.12146

Froese R, Pauly D (Eds.) (2025). FishBase: Encrasicholina heteroloba (Bleeker, 1852). Retrieved November 3, 2025, from https://www.fishbase.org/summary/Encrasicholina-heteroloba.html

Fujimoto, M (2018) Economic impact of the dagaa processing industry on a coastal village in Zanzibar, Tanzania. African Study Monographs, Supplement 55: 145–162

Groeneveld JC, K.A. Koranteng (2016) The RV Dr Fridtjof Nansen in the Western Indian Ocean: voyages of marine research and capacity development, FAO, Rome, Italy.

Gulland JA (1971) The fish resources of the ocean. FAO Fish. Tech. Pap. openknowledge.fao.org/handle/20.500.14283/al937e

Harlyan LI, Wu D, Kinashi R, Kaewnern M, Matsuishi T (2019) Validation of a feedback harvest control rule in data-limited conditions for managing multispecies fisheries. Can. J. of Fish. Aquat. Sci. 76(10): 1885–1893. 10.1139/cjfas-2018-0318

He JX, Stewart DJ, Rudstam LG (2016) Growth parameters as growth indices in time-varying environments: a comparison among four approaches to using the von Bertalanffy growth function. Oneida Lake: long-term dynamics of a managed ecosystem and its fishery. Am. Fish. Soc. Bethesda, MD. 2016:475–96.

Herrón P, Mildenberger TK, Díaz JM, Wolff M (2018) Assessment of the stock status of small-scale and multi-gear fisheries resources in the tropical Eastern Pacific region. Reg. Stud. Mar. Sc. 24: 311–323. 10.1016/j.rsma.2018.09.008.

Horbowy J, Hommik K (2022) Analysis of *F*_msy_ in light of life-history traits. Effects on its proxies and length-based indicators. Fish Fish. 23: 663–679. 10.1111/faf.12640

Hordyk AR, Ono K, Valencia SR, Loneragan NR, Prince JD (2015) A novel length-based empirical estimation method of spawning potential ratio (SPR), and tests of its performance for small-scale, data-poor fisheries. ICES J. Mar. Sc. 72(1): 217–231. 10.1093/icesjms/fsu004

Ibengwe LJ, Onyango PO, Hepelwa AS, Chegere MJ (2022) Regional trade integration and its relation to income and inequalities among Tanzanian marine dagaa fishers, processors and traders. Mar. Policy. 137: 104975. 10.1016/j.marpol.2022.104975.

Ibengwe LJ, Onyango PO, Hepelwa AS, Mfilinge PL (2023) Revealing the hidden marine *dagaa* cross-border trade in mainland Tanzania. Rev Fish Biol Fisheries 33: 717–738. 10.1007/s11160-023-09769-4

Ingles J, Pauly D (1984) An atlas of the growth, mortality and recruitment of Philippines fishes. ICLARM Tech. Rep. 13: 127.

Jebri F, Z.L. Jacobs, D.E. Raitsos, M. Srokosz, S.C. Painter, S. Kelly, M.J. Roberts, L. Scott, S.F.W. Taylor, M. Palmer, H. Kizenga, Y. Shaghude, J. Wihsgott, E. Popova (2020) Interannual monsoon wind variability as a key driver of East African small pelagic fisheries, Nature Publishing Group, UK. 10.1038/s41598-020-70275-9.

Jones R (1984) Assessing the effects of changes in exploitation pattern using length composition data (with notes on VPA and cohort analysis). FAO Fish. Tech. Pap. 256.

Juliani AS, Saputra SW, Helminuddin (2019) Sustainability assessment of Devis’ anchovy (Encrasicholina devisi (Whitley, 1940)) (Clupeiformes: Engraulidae) fisheries based on biology aspects, Kutai Kartanegara, Indonesia. AACL Bioflux 12(5):1938–1950.

Kamukuru AT, Mahongo SB, Sekadende BC, Sululu, JS (2020) Age, growth and mortality of the anchovy Stolephorus commersonnii (Lacepède, 1803) (Clupeiformes) caught off the coast of Tanga, Tanzania. Western Indian Ocean Journal of Marine Science, 1/2020, 95–104. 10.4314/wiojms.si2020.1.9

Lavapie-Gonzales F, Ganaden SR, Gayanilo FC (1997) Some population parameters of commercially important fishes in the Philippines. Bureau of Fisheries and Aquatic Resources, Philippines. 114.

Losse GF (1966) Fishes taken by purse-seine and dipnet in the Zanzibar Channel, East Afr. Agric. For. J. 32: 50–53. 10.1080/00128325.1966.11662091.

Mace PM (2001) A new role for MSY in single-species and ecosystem approaches to fisheries stock assessment and management. Fish Fish. 2: 2–32, 10.1046/j.1467-2979.2001.00033.x

Matsuishi TF (2025) Status of Southeast Asian fisheries: distinctive characteristics and pathways to sustainable fisheries. Fish. Sci. 91: 205–216. 10.1007/s12562-025-01854-w

Mayala P, D.M.K. Mayala (2016) Assessment of socio-economic value of the small pelagic fishery in Mafia Island, United Nations University Fisheries Training Programme, Iceland, Tanzania.

Mildenberger TK, Taylor MH, Wolff M (2017) TropFishR: an R package for fisheries analysis with length-frequency data. Methods Ecol. Evol. 10.1111/2041-210X.12791

MLF (2015) The United Republic of Tanzania Ministry of Livestock and Fisheries Development National Fisheries Policy, 2015, 2015. https://www.mifugouvuvi.go.tz/uploads/publications/en1595837773-NATIONAL%20FISHERIES%20POLICY%202015.pdf

MLF (2016) The United Republic of Tanzania, Ministry of Livestock and Fisheries: Marine Fisheries Frame Survey 2016 Report Mainland Tanzania, 2016.

Muhando CA, Chikambi K, Rumisha (2008) Distribution and status of coastal habitats and resources in Tanzania, Report submitted to WWF – Dar es Salaam. https://scholar.google.com/scholar?hl=en&as_sdt=0%2C5&q=C.A.+Muhando+DISTRIBUTION+AND+STATUS+OF+COASTAL+HABITATS+AND+RESOURCES+IN+TANZANIA&btnG=

Muller RG (1976) Population biology of *Stolephorus heterolobus* (Pisces: Engraulidae) in Palau, Western Caroline Islands. Ph.D. thesis, University of Hawaii. 174.

Musel J, Anuar A, Hassan MH, Mustafa WZW, Paul PS, Sahidun I, Chiba SUA (2022) Population dynamics and the spawning season of the commercial dominant species (*Encrasicholina devisi* and *Sardinella fimbriata*) from the northern region of Sarawak. Aquac. Aquar. Conserv. Legis. 15(3): 1162–1177.

Nhwani LB, E.D.M. Nhwani (1988) Aspects of the fishery biology of small pelagic fishes at Dar Es Salaam, Tanzan., Fishbyte: 7–10.

Padilla JE (1991) Managing tropical multispecies fisheries with multiple objectives. Ph.D. thesis, Simon Fraser University. 235.

Paula e Solver R (1992) Growth of the buccaneer anchovy *Encrasicholina punctifer* off Mozambique, based on samples collected in research surveys. Rev. Invest. Pesq. (Maputo) 21: 69–78.

Pauly D (1978) A preliminary compilation of fish length growth parameters. Ber. Inst. Meereskd. Christian-Albrechts-Univ. Kiel (55):1–200.

Pauly D (1980) On the interrelationships between natural mortality, growth parameters, and mean environmental temperature in 175 fish stocks. ICES J. Mar. Sci. 10.1093/icesjms/39.2.175

Pauly D (1990) Length-converted catch curves and the seasonal growth of fishes. Fishbyte. 8:33–38.

Pauly D, Munro JL (1984) “Once more on the comparison of growth in fish and invertebrates.” Fishbyte 2: 21.

Pauly D, Moreau J, Abad N (1995) Comparison of age-structured and length-converted catch curves of brown trout *Salmo trutta* in two French rivers. Fish Res. 22:197–204. 10.1016/0165-7836(94)00323-O.

Peter HK, Kuguru B, Sailale I, Marwa G, Semba M (2023) Spatial and temporal distribution of small pelagic fishes in the territorial waters of Tanzania. Version 1.9. TanBIF. Occurrence dataset. 10.15468/d5jjq2. Accessed via GBIF.org on 2025-04-24.

Prince JD, Hordyk A, Valencia SV, Loneragan N, Sainsbury K (2015) Revisiting the concept of Beverton–Holt life history invariants with the aim of informing data-poor fisheries assessment. ICES J. Mar. Sc. 72: 194–203. 10.1093/icesjms/fsu011

Rawlinson N (1989) Population dynamics of the commercially important baitfish species *Stolephorus heterolobus* in Solomon Islands. Fishbyte. 7(1): 12–17.

Raymond JHB, Hylen A, Østvedt OJ, Alvsvaag J, Iles TC (2004) Growth, maturation, and longevity of maturation cohorts of Norwegian spring-spawning herring, ICES J. Mar. Sci. 61(2): 165–175. 10.1016/j.icesjms.2004.01.001

Sekadende B, Scott L, Anderson J, Aswani S, Francis J, Jacobs Z, Jebri F, Jiddawi N, Kamukuru AT, Kelly S, Kizenga H, Kuguru B, Kyewalyanga M, Noyon M, Nyandwi N, Painter. SC, Palmer M, Raitsos DE, Roberts M, Sailley SF, Samoilys M, Sauer WHH, Shayo S, Shaghude Y, Taylor SFW, Wihsgott J, Popova E (2020) The small pelagic fishery of the Pemba Channel, Tanzania: what we know and what we need to know for management under climate change, Ocean Coast. Manag. 197 (2020), 10.1016/j.ocecoaman.2020.105322

Sparre P, Venema SCW (1998) Introduction to tropical fish stock assessment. Part 1. Manual. FAO Fish Tech Paper. 306: 1–407.

Supeni EA, Dobo J (2017) Exploitation status of Devi’s anchovy in Kei Island waters: based on total length data. IOCP Conf. Series: Earth and Environmental Science. 89: 012004. doi:10.1088/1755-1315/89/1/012004

Tanaka S (1960) Studies on the dynamics and management of fish populations. Bull. Tokai. Reg. Fish. Res. Lab., 28: 1–200.

Taufani WT, Matsuishi TF (2024) Precision of estimated growth parameters of yellowfin tuna (*Thunnus albacares*) from length-frequency data estimated by bootstrapping. Fish. Manage. Ecol. 10.1111/fme.12781

Tham AK (1967) A contribution to the study of the growth of members of the genus *Stolephorus* Lacepede in Singapore Strait. Proc. IPFC. 12(2): 1–25.

Thordarson G, Björnsson H, Hjörleifsson E, Steinarsson BÆ (2006) Are Fmax and F0.1 Illusive as Fisheries Reference Points? ICES NWWG WD, 30

Thompson WF, Bell FH (1934) Biological statistics of the Pacific halibut fishery 2. Effect of changes in intensity upon total yield and yield per unit of gear. Rep. Int. Fish. Commun. 8:1–49.

von Bertalanffy L (1938) A quantitative theory of organic growth (inquiries on growth laws. II). Human Biology 10(2):181–213

Wang Q, Yang JX, Zhou GQ, Zhu YA, Shan H (2011) Length–weight and chelae length–width relationships of the crayfish *Procambarus clarkii* under culture conditions. J. Freshw. Ecol, 26:2, 287–294, Doi:10.1080/02705060.2011.564380

Wang K, Zhang C, Xu B, Xue Y, Ren Y (2020) Selecting optimal bin size to account for growth variability in Electronic LEngth Frequency ANalysis (ELEFAN). Fish. Res. 10.1016/j.fishres.2019.105474

